# A Smart Region-Growing algorithm for single-neuron segmentation from confocal and 2-photon datasets

**DOI:** 10.1101/287029

**Authors:** Alejandro Luis Callara, Chiara Magliaro, Arti Ahluwalia, Nicola Vanello

## Abstract

Accurately digitizing the brain at the micro-scale is crucial for investigating brain structure-function relationships and documenting morphological alterations due to neuropathies. Here we present a new Smart Region Growing algorithm (SmRG) for the segmentation of single neurons in their intricate 3D arrangement within the brain. Its Region Growing procedure is based on a homogeneity predicate determined by describing the pixel intensity statistics of confocal acquisitions with a mixture model, enabling an accurate reconstruction of complex 3D cellular structures from high-resolution images of neural tissue. The algorithm’s outcome is a 3D matrix of logical values identifying the voxels belonging to the segmented structure, thus providing additional useful volumetric information on neurons.

To highlight the algorithm’s full potential, we compared its performance in terms of accuracy, reproducibility, precision and robustness of 3D neuron reconstructions based on microscopic data from different brain locations and imaging protocols against both manual and state-of-the-art reconstruction tools.

## 1. INTRODUCTION

Digitalizing a high-fidelity map of the neurons populating the brain is a central endeavour for neuroscience research, and a crucial step for the delineation of the full Connectome (Alivisatos et al. 2012). Moreover, single-neuron reconstruction from empirical data can be used to generate models and make predictions about higher-level brain organization, as well as to study the normal development of dendritic and axonal arbours or document neuro-(patho)physiological changes (Budd et al. 2015).

Doubtless, confocal and two-photon microscopy are the best candidates to image defined cellular populations in three-dimensional (3D) biological specimens (Ntziachristos 2010; Wilt et al. 2009). Their imaging depth, as well as the quality of the acquired datasets can be further improved thanks to recent tissue-clearing solutions, which render brain tissue transparent to photons by reducing the source of scattering, allowing confocal acquisitions with enhanced Signal to Noise Ratios and Contrast to Noise Ratios while maintaining low laser power (Chung and Deisseroth 2013; Chiara Magliaro et al. 2016; Richardson and Lichtman 2015). While these emerging technologies and protocols, combined with fluorescence-based labelling techniques, enable the imaging of the brain’s intricacies at the microscale, single-cell segmentation algorithms able to deal with these datasets are still lacking, despite targeted initiatives such as the DIADEM (DIgital reconstructions of Axonal and DEndrite Morphology) challenge in 2009-2010 (Gillette et al. 2011) and the BigNeuron project in 2015 (Peng et al. 2015). Indeed, many tools and algorithms for neuron segmentation primarily focus on sparsely labelled data, such that their application to images (or volumes) representing densely packed neurons, typical of mammalian brains, is limited (Chothani et al. 2011; Hernandez et al. 2018; Peng et al. 2014; C.-W. Wang et al. 2017; Y. Wang et al. 2011). As a matter of fact, most of the state-of-the-art algorithms are purposely developed to deal with low-quality images in which there may be (i) noisy points causing over-tracing, (ii) gaps between continuous arbours causing under-tracing and (iii) non-smooth surfaces of the arbours violating geometric assumptions (Liu et al. 2016).

Confocal and 2-photon datasets are characterized by on-plane and intra-plane pixel intensity heterogeneities, deriving from optical phenomena and the non-uniform distribution of fluorophores through the sample (Diaspro 2001). Given these intrinsic features, a valid procedure for accurately digitizing the neural structures in the stack could be obtained by leveraging on local approaches and methods enforcing spatial constraints, such as region growing procedures (RG) (Acciai et al. 2016; Brice and Fennema 1970; Xiao and Peng 2013). RG is a pixel intensity-based segmentation method that identifies a Region of Interest (ROI) starting from a pixel, i.e. the seed, belonging to the ROI itself. The neighbouring pixels of the seed are iteratively examined based on a predefined rule, usually a homogeneity predicate, which can be estimated locally to determine whether they should be added to the ROI or not. The performance of the procedure may be influenced by both the seed selection and the rule (Baswaraj et al. 2012). The choice of the rule may be nontrivial, in particular in view of delivering a general-purpose segmentation algorithm. Here we propose a novel RG strategy based on an estimation which considers the image formation process to define intrinsic properties of signal distribution in the image in question.

Our rationale is that confocal and 2-photon microscopy are based on sampling successive points in a focal plane to reproduce the spatial distribution of fluorescent probes within a sample. Hence, each pixel contains a discrete measure of the detected fluorescence within a sample interval, represented by a photon count, and certain amount of noise, deriving from different sources (Calapez and Rosa 2010; Pawley and Pawley 2006). Therefore, statistical methods represent a natural way of describing confocal or 2-photon datasets. Different models have been proposed to depict confocal image properties (Calapez et al. 2002; Pawley and Pawley 2006). Specifically, mixture models (MM) have been suggested as the best descriptor of the sharp peaks and the long tails typical of background and low fluorescence distributions (Calapez and Rosa 2010).

Given these considerations, we have developed a new Smart Region Growing algorithm (SmRG), which couples the RG procedure with a MM describing the signal statistics, to calculate local homogeneity predicates (i.e. local thresholds) for iteratively growing the structure to be segmented. Here, we describe the SmRG workflow for single-neuron segmentation. Then, we evaluate its performance in segmenting different neuron types from confocal and 2-photon datasets, comparing the results with those obtained with a gold standard manual reconstruction. Furthermore, we compare our algorithm with state-of-the-art (SoA) tools widely used in the field of neuron reconstruction.

## 2. SMART REGION GROWING ALGORITHM (SmRG): AN OUTLINE

The SmRG is an open-source algorithm developed in Matlab (The Mathworks-Inc, USA). A package of the functions needed for running the algorithm are available at http://www.centropiaggio.unipi.it/smrg-algorithm-smart-region-growing-3d-neuron-segmentation.

The SmRG is driven by a homogeneity predicate for establishing a local threshold based on the intensity levels of confocal datasets. Specifically, it exploits the statistics of the background and the signal distributions of the confocal acquisitions and a linear MM to determine the probability with which a given pixel (voxel) can be considered as part of the ROI or not (see Appendix I for details about the MM). The rule to grow regions is then designed from these probabilities.

The workflow of the SmRG is sketched in Fig 1. It begins by selecting a seed, either manually or automatically (Fig 1A).

**Fig 1.**
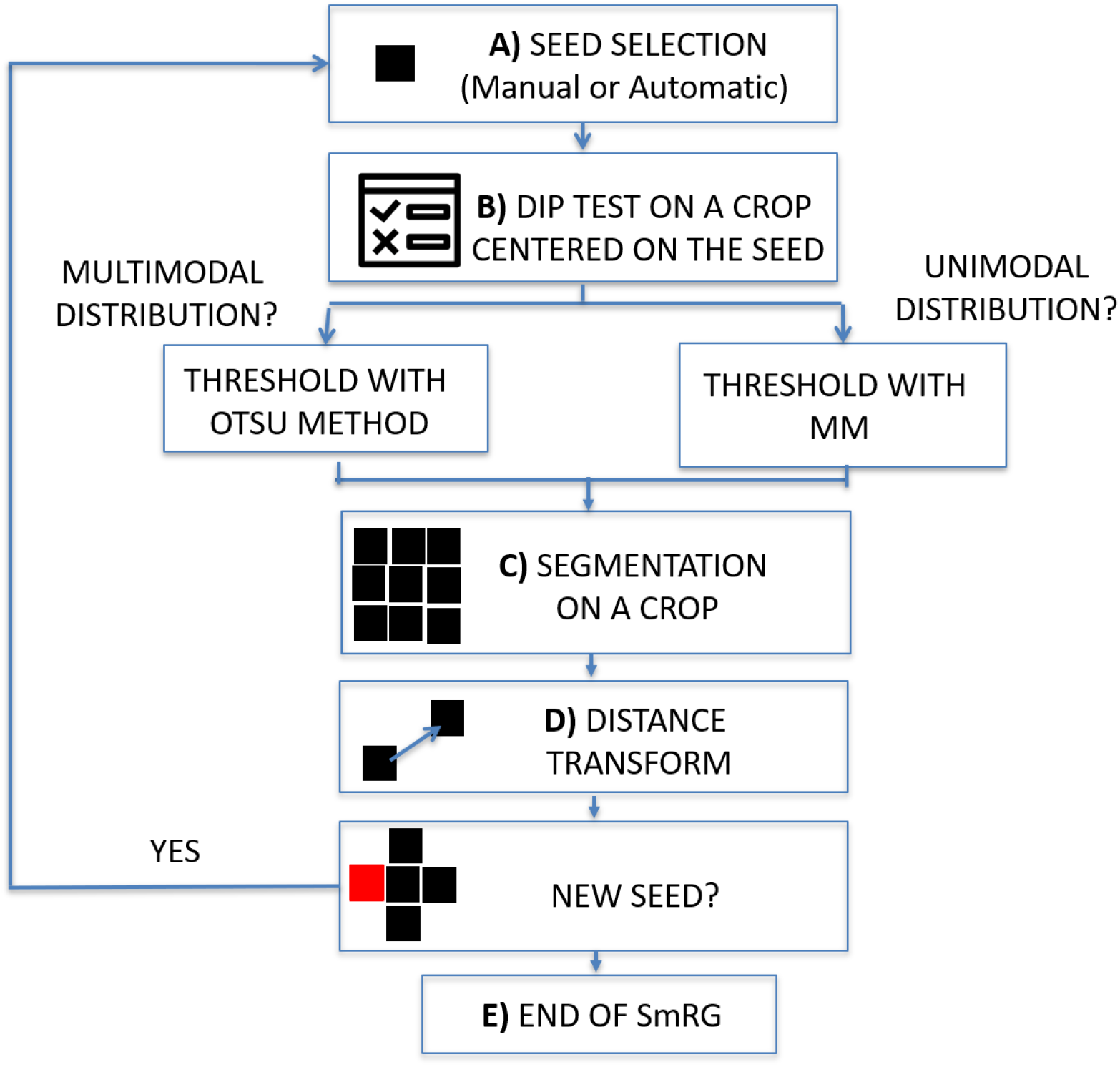
Workflow of the SmRG. A) Manual or automatic seed selection. B) Dip test to test for unimodality against multimodality on a MxNx3 crop centred on the seed. The threshold is determined with Otsu’s method or through the Mixture Model according to whether the distribution is multimodal or not. C) 3D segmentation of a MxNx3 crop. D) The regional maxima of the distance transform of the segmented MxNx3 crop are chosen as new seeds. E) The procedure iterates until there are no more new seeds.

In the first case, the user is asked to identify the seed position by selecting a point on a focal plane (e.g. a pixel belonging to the soma), while in the latter the Hough transform (Nixon and Aguado 2012) searches for spherical objects within the stack to identify the somata: the seed (or the seeds) is (or are) chosen as the centre of the detected sphere (or spheres). Then, the homogeneity predicate is derived locally on an image volume centred on the seed. The volume dimension is a trade-off between the goodness-of-fit of the MM and the localness of the segmentation, and by default is set 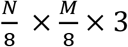, where N and M are the on-plane size of the image stack. To ensure enough data points for MM fitting, the crop size is never smaller than 32×32×3. At this step a Hartigan’s dip test (Hartigan and Hartigan 1986) (p<0.01) is performed on the pixel intensity distribution of the crop to test for unimodality against multimodality (Fig 1B). In the case of multimodality the segmentation proceeds with Otsu’s method (Otsu 1979), a well-known thresholding technique for multimodal distributions (Guo et al. 2012). Otherwise, a linear MM, considering the background as a normal distribution and the signal as a negative binomial, is fitted by means of an Expectation Maximization (EM) algorithm on the crop pixel intensity distribution. Indeed, mixture models combining normal and negative binomial distributions have been observed to fully characterize the signal associated with confocal images (Calapez and Rosa 2010) (see Appendix 1 for details about the MM and model fitting). The homogeneity predicate is derived from the posterior probability of the MM, *α (or 1−α)*, denoting the probability at which a given pixel can be considered as part of the background (or the signal) distribution. The rule is thus obtained as user defined threshold for *α* (e.g. with *1−α>0.999* all the seed’s neighbouring pixels whose probability of belonging to the signal exceeds 99.9% are segmented) (Fig 1C). Each pixel that satisfies this rule and is spatially connected to the seed within the crop is added to the ROI. At this point, new seeds are chosen from the points just recognized as part of the neuron to be segmented. In particular, for each segmented plane the regional maxima of the distance transform (Maurer and Raghavan 2003) are taken as new seeds (Fig 1D). The algorithm iterates for each detected seed and the process stops when there are no more pixels to add (Fig 1E).

The result of the SmRG is a 3D matrix of logical values, whose true values (i.e. logical ones) represent the voxels constituting an isolated neuron. Fig 2 shows an example of a Purkinje cells segmented using the SmRG from a confocal dataset representing a 1mm-thick slice from murine cerebellum, obtained after applying the CLARITY protocol described in Magliaro et al., 2016(C. Magliaro et al. 2016).

**Fig 2.**
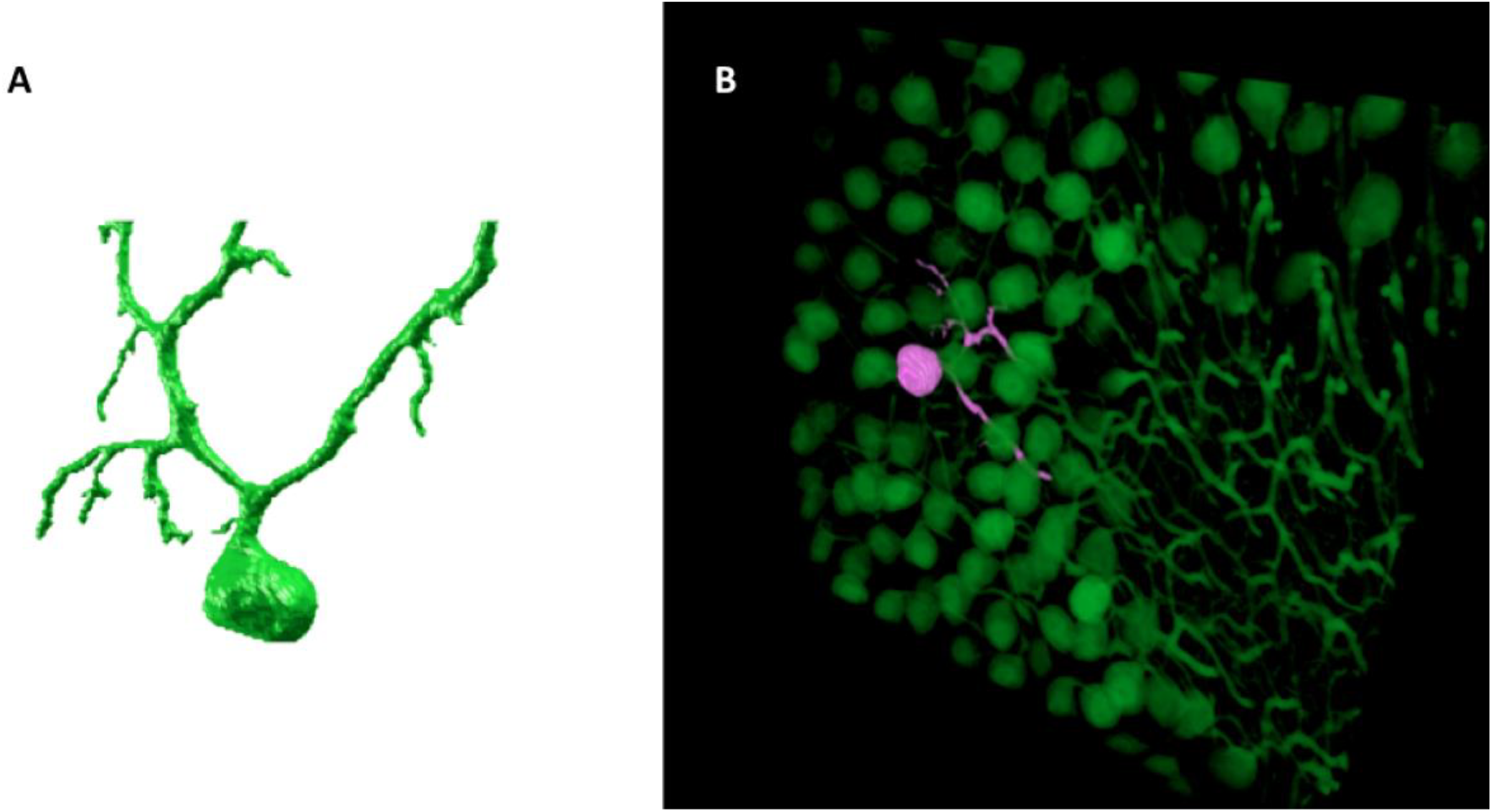
An example of SmRG outcome: A) a Purkinje cell from clarified murine cerebellum acquired using a Nikon A1 confocal microscope; B) the same Purkinje cells identified within its confocal dataset.

## 3. MATERIALS AND METHODS

To evaluate the SmRG’s performance, we processed two different sets of data. First, confocal acquisitions of 1 mm-thick slices of clarified cerebellum from a L7GFP mouse were analysed to isolate Purkinje Cells (PCs) expressing Green Fluorescent Protein (GFP). The aim was to demonstrate i) the SmRG’s accuracy with respect to a manual segmentation performed by experts, as it is still considered the gold standard for neuron segmentation (Al-Kofahi et al. 2003; Meijering 2010), ii) the SmRG’s reproducibility and iii) its ability to handle 3D microscopic datasets representing dense-packed neurons compared with other tools available in literature.

Then, Olfactory Projection (OP) Fibers dataset from the DIADEM challenge was processed with the SmRG. The SmRG reconstructions were quantitatively compared to the manually-traced gold-standards provided by the DIADEM. Moreover, 3D neuron segmentation was performed using other SoA tools evaluating the outputs against the DIADEM gold standards through the metrics SD, SSD and SSD%. This allowed an assessment of the SmRG’s ability to reconstruct 3D neuron morphology with the same precision as SoA algorithms.

The tools used for both PC and OP datasets were the Vaa3D (version 3.200) app2 (Xiao and Peng 2013), MST-tracing (Basu and Racoceanu 2014), SIGEN (Ikeno et al. 2018) and MOST (Ming et al. 2013) plug-ins. They have been extensively validated in other reports and are widely used to compare reconstructions provided by new segmentation algorithms (Liu et al. 2016; Peng et al. 2014).

### 3.1 Datasets representing PCs

#### 3.1.1 Accuracy test: SmRG algorithm versus manual segmentation

The confocal datasets representing dense-packed PCs from 1 mm-thick slices from clarified L7GFP murine cerebellum were those already manually segmented in Magliaro et al., 2017 (Chiara Magliaro et al. 2017). They are available for download at http://www.centropiaggio.unipi.it/mansegtool. Specifically, n = 3 Purkinje cells were segmented automatically with the SmRG algorithm and manually by 6 experts with the ManSegTool, a tool purposely developed for facilitating the manual segmentation of 3D stacks ^35^.

The SmRG’s segmentation accuracy was evaluated by comparing morphometric features extracted from the two outputs. Briefly, we considered i) the surface area, ii) the volume and iii) the Sholl analysis (Chiara Magliaro et al. 2017; Sholl 1955) of segmented structures. To compare Sholl profiles, we calculated the total area under the curve (AUC) using the trapezoidal rule thus obtaining a single measure for each profile (Binley et al. 2014). Statistical differences between the features in the manual segmented structures and those resulting from the SmRG were evaluated by means of the Friedman’s test with replicates. Friedman’s test allows testing treatments under study (i.e., columns) after adjusting for nuisance effects (i.e., rows). Replicates refer to more than one observation for each combination of factors. In our case, surface area, volume and the AUC of Sholl profiles were blocking factors (i.e., rows) with replicates represented by the three neurons, while users and SmRG represented treatments (i.e., columns). Thus, we are testing the null hypothesis of no difference between manual and SmRG-based segmentation.

#### 3.1.2 SmRG reproducibility

Reproducibility tests were performed by segmenting the same n=3 PCs starting from different seeds. Specifically, we randomly chose 10 pixels picked from different regions of the neuron. Volume, surface area and AUC of Sholl profiles were obtained for each seed and the reproducibility was quantified for each neuron as the coefficient of variation of each measure (i.e. the standard deviation normalized by the mean).

#### 3.1.3 SmRG vs SoA tools

In order to highlight the SmRG’s ability to segment single-neurons from confocal datasets represented densely-packed cells, we processed a 3D image stack with the App2, MST, SIGEN and MOST Vaa3d plugins. Reconstructions provided by these tools and by SmRG were visually compared.

### 3.2 DIADEM datasets representing OP Fibers

The dataset representing OP Fibers is available at http://diademchallenge.org/olfactory_projection_fibers_readme.html. It contains 9 separate drosophila olfactory axonal projection image stacks acquired with a two-photon microscope and their respective gold standard reconstructions provided by the DIADEM (Evers et al. 2005; Jefferis et al. 2007). We segmented all the neurons except OP2, since it contains many irrelevant structures (Liu et al. 2016). The SmRG and SoA algorithm reconstructions were compared with the DIADEM gold-standards. Comparisons between automatic tools were made by means of the following metrics: i) the spatial distance (SD) ii) the substantial spatial distance (SSD) and iii) the percentual substantial spatial distance (%SSD). The spatial distance is estimated as it follows:

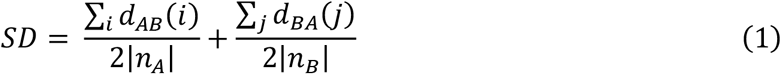

With

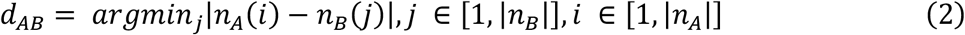

and

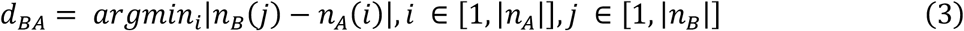

i.e., given two reconstructions, A and B, the spatial distance is obtained by averaging the Euclidean distance between the nodes of A and the nodes of B, i.e. *d*_*AB*_, with the reciprocal measure, i.e. *d*_*BA*_. Specifically, *d*_*AB*_ is obtained by selecting, for each node belonging to A, the minimum among the distances with each other node of B. *d*_*AB*_ is thus obtained by repeating this operation for each node of A, and then by averaging the results. The same operation is performed with the nodes belonging to B, to obtain *d*_*BA*_.

The SSD is obtained by selecting the node pairs in A and B with a minimal distance above a given threshold S, and then performing their average. Specifically, given:

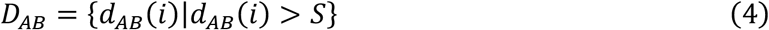

and

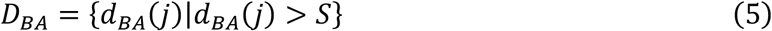

Then, the SSD is defined as follows:

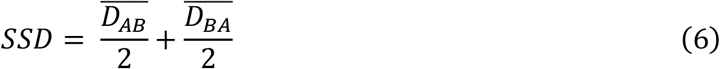

Finally, the %SSD is obtained by estimating the fraction of nodes contributing to SSD. These metrics express the similarity of two different reconstructions (Peng, Long, Zhao, et al. 2011). Essentially, SD is a measure of how different 2 reconstructions are, while SSD and SSD% measure the extent of differences between 2 reconstructions considering only points above a tolerance threshold S. The tolerance threshold for the evaluation of the SSD metric was 2 (i.e. S=2) voxels, as suggested in Peng et al., 2011(Peng, Long, and Myers 2011). Given that the SmRG’s output is a 3D logical matrix constituting the whole neuron, while the DIADEM gold-standard is a set of points of interest (i.e. a *.swc file), a thinning procedure was necessary to reduce the volumetric information to a skeleton. To this end, we calculated the 3D skeleton of the SmRG output via a 3-D Medial Surface Axis Thinning Algorithm (Lee et al. 1994). From the points constituting the skeleton we reconstructed the corresponding *.swc file using an *ad hoc* routine, ensuring a fair mapping between the DIADEM reference points and the SmRG ones.

Moreover, the precision, recall and F-score of the SmRG reconstructions were determined with respect to the gold-standard, quantifying the spatial overlap between the closest corresponding nodes of the two reconstructions (Powers 2011) and varying the tolerance threshold from 2 to 5 voxels, to evaluate the SmRG’s sensitivity to this parameter (Radojević and Meijering 2018).

## 4. RESULTS

### 4.1 Purkinje Cell segmentation

#### 4.1.1 SmRG vs manual segmentation

Fig 3 shows an example of the same PC segmented by an expert and by the SmRG.

**Fig 3.**
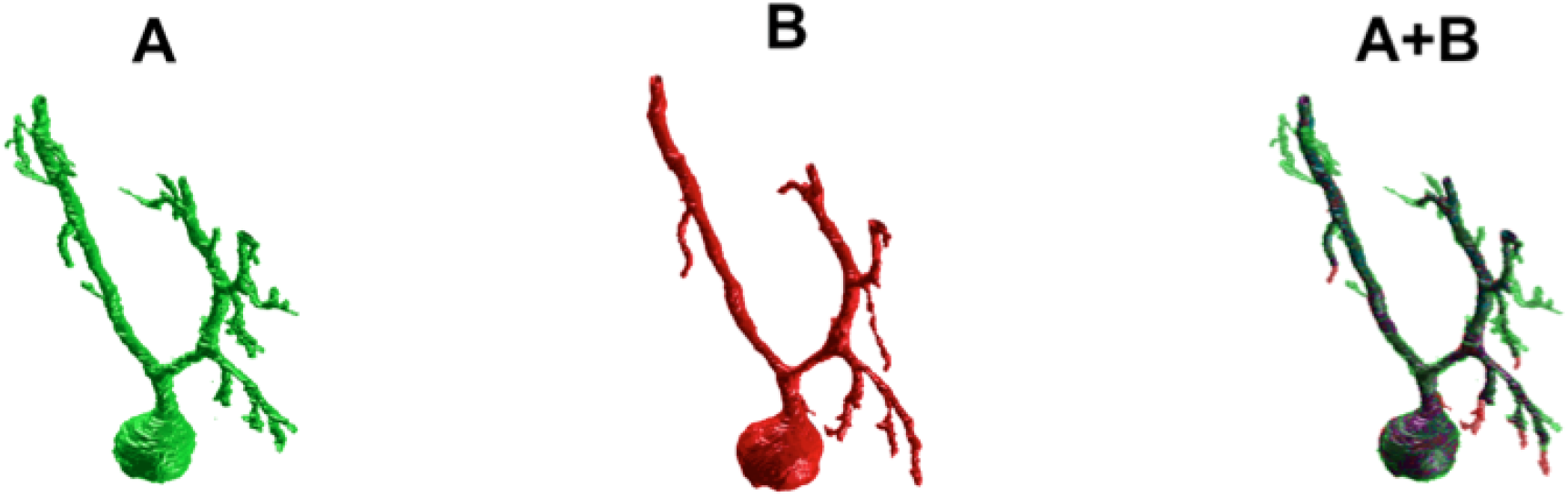
SmRG versus Manual Segmentation. A) Gold-standard manual segmentation. B) SmRG automatic segmentation. C) Merge of manual (green) and automatic (red) segmentation, common voxels are reported in purple.

The SmRG’s accuracy was assessed by comparing volume, surface area and AUC of Sholl profiles extracted from the segmented PCs with the results obtained by manually segmented ones (Fig 4).

**Fig 4.**
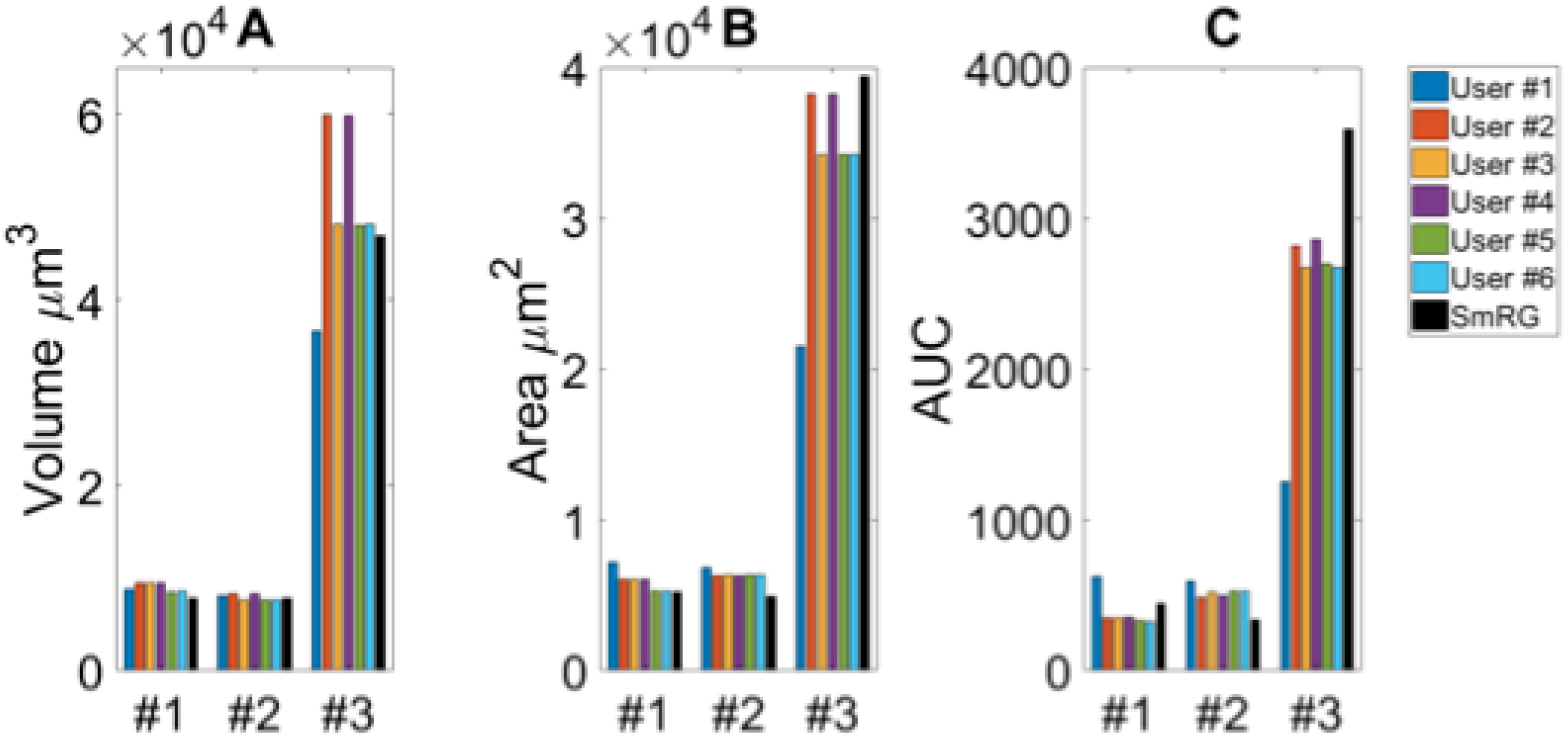
Testing SmRG accuracy. A) Neuron volume. B) Neuron surface. C) AUC (area under the curve) of Sholl profiles. Friedman’s test was performed with Volume, Area and AUC as blocking factors (rows, nuisance effects) with replicates (neurons #1, #2 and #3), and with users and SmRG as treatments (column). No statistical differences were observed (p-value=0.8233).

The single-neuron reconstructions provide quantitative information on the morphology of individual neurons in their native context where they are surrounded by neighbouring cells. Clearly the algorithm developed is able to follow neurite arborisation, segmenting smaller branches with similar performance to manual segmentation. Furthermore, the structure obtained with the SmRG is consistently characterized by a smooth volume, compared with the manual segmentation. A typical example is reported in Fig 5, showing a zoomed detail of manual and SmRG segmentation results.

**Fig 5.**
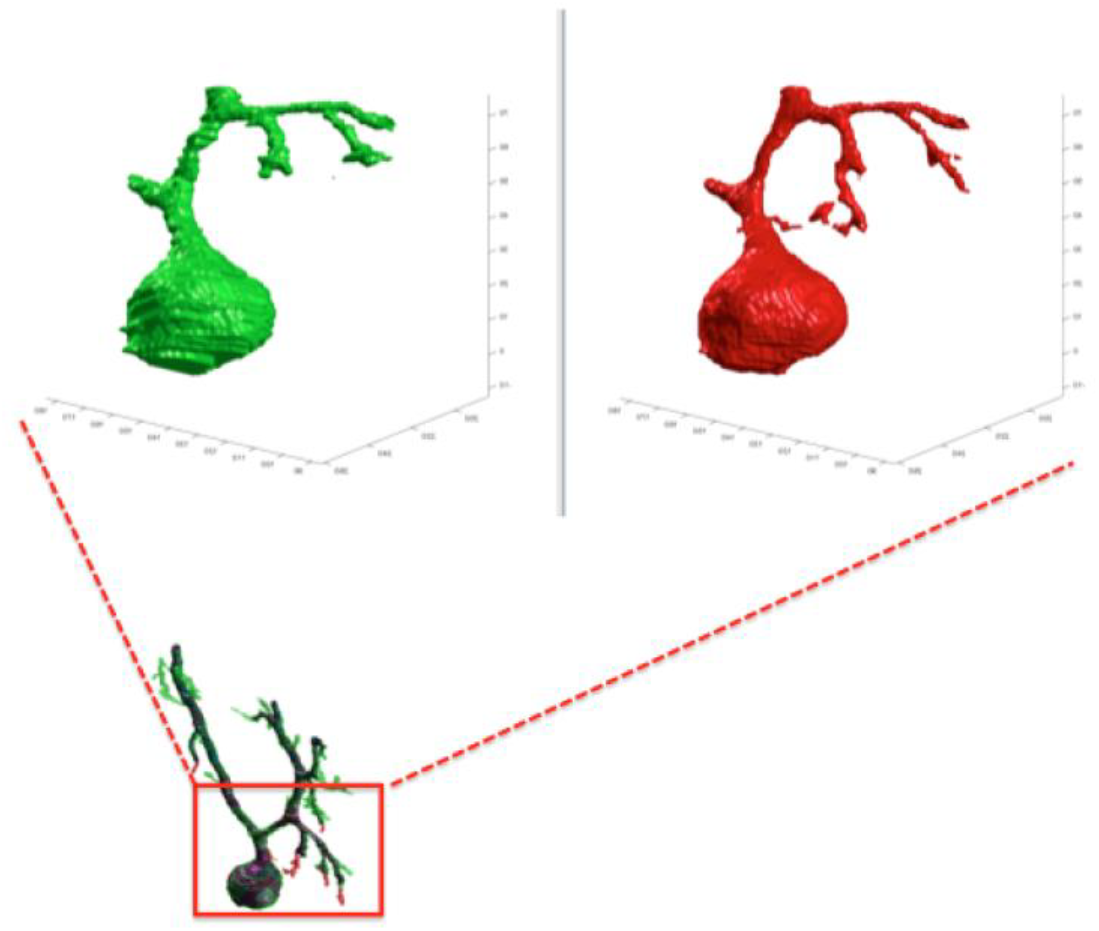
A detail of the manual and SmRG neuron reconstruction. It is clear that the SmRG segmentation (red) leads to a smoother volume than the manual (green) one.

The Friedman’s test showed no significant differences between the SmRG and the ManSegTool segmentation in terms of surface area, volume and Sholl profiles of the segmented structures (p = 0.8233); a detailed ANOVA table of the Friedman’s test is reported in Table 1. In summary, the results in the table demonstrate that the SmRG’s performance is comparable to that obtained from manual segmentation performed by experts in terms of the accuracy of the morphological parameters considered.

**Table 1.** Friedman’s ANOVA table. SS=Sum of Squares due to each sources; Df = Degree of freedom associated with each source; MS = Mean Squares, which is SS/Df; Chi-sq: Freedman Chi-square statistic; p: p-value for the Chi-square statistic.

| Source | SS | Df | MS | Chi-sq | p>Chi-sq |
| --- | --- | --- | --- | --- | --- |
| Columns | 110.94 | 6 | 10.4907 | 2.08 | 0.8233 |
| Interaction | 64.89 | 12 | 5.4074 |  |  |
| Error | 2132.67 | 42 | 50.7778 |  |  |
| Total | 2308.5 | 62 |  |  |  |

#### 4.1.2 SmRG’s reproducibility

Table 2 reports the coefficients of variation of volume, surface area and AUC of Sholl profiles for each segmented neuron. The maximum coefficient of variation was equal to 0.0258, demonstrating the robustness of the SmRG to changes in initial conditions (i.e. the position of a seed belonging to the structure of interest).

**Table 2.**
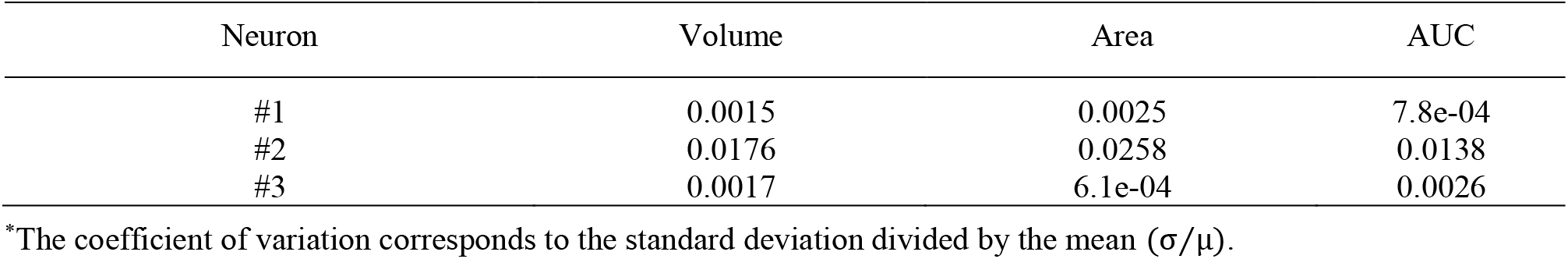
Results of SmRG’s reproducibility. Coefficients of variation for neuron volume, area and AUC for n different RG seeds.

#### 4.1.3 SmRG vs other tools

Fig 5 shows an example of the outputs obtained segmenting the same confocal 3D stack with the App2, MST, SIGEN and MOST routines and with the SmRG. We were only able to assess the comparisons visually, since none of Vaa3D plugins was able to handle such dense datasets.

### 4.2 OP fibers: SmRG vs the DIADEM gold-standard

OP fibers segmented with the SmRG are reported in Fig 6, along with the manually-traced gold-standard provided by the DIADEM.

**Fig 6.**
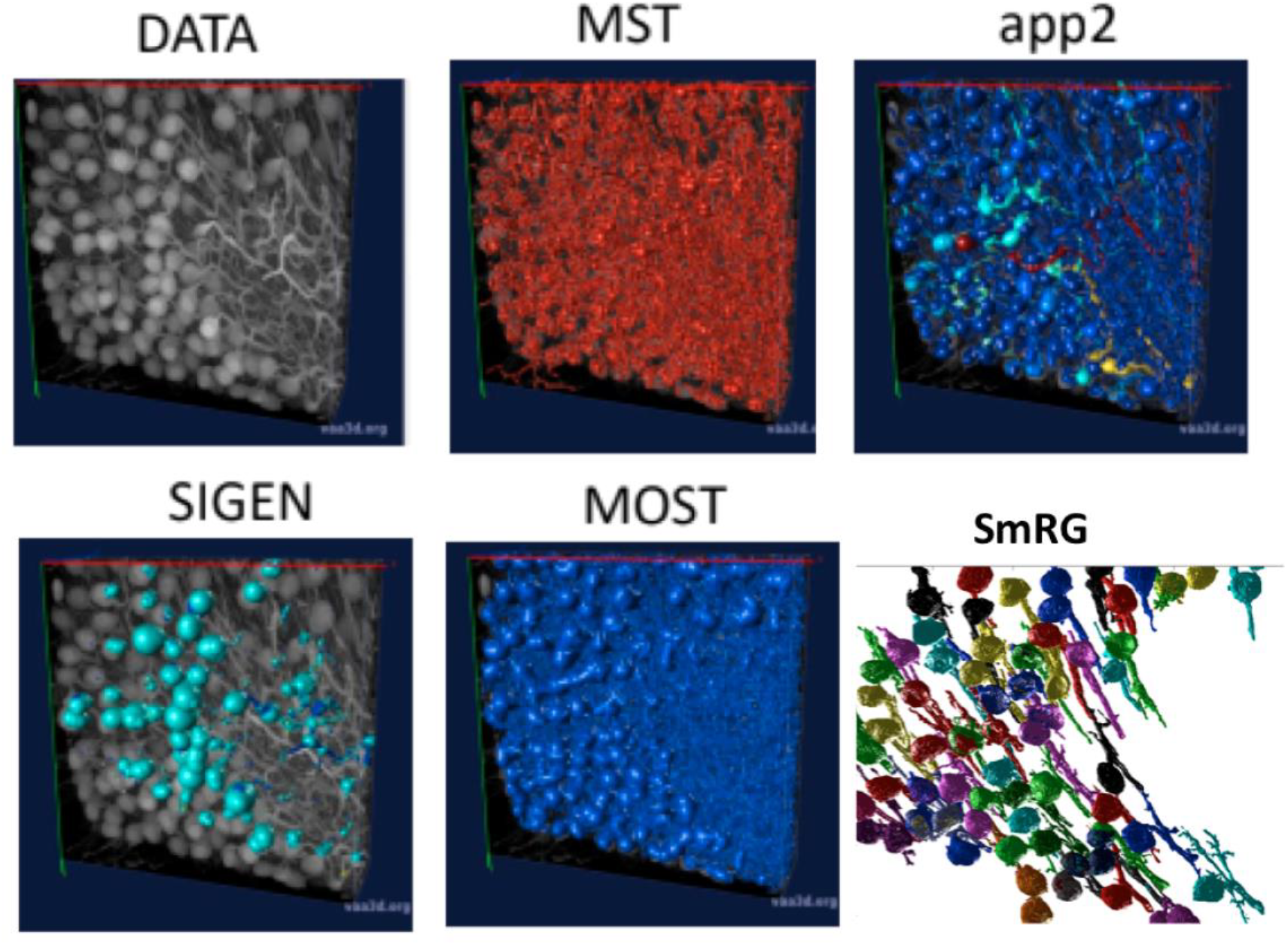
An example of a confocal dataset representing PCs from a clarified L7GFP murine cerebellum, segmented with MST, app2, MOST, SIGEN and SmRG. None of the SOA tools is able to deal with this dense dataset, while the SmRG is able to isolate the PCs within the dataset. Different colours refer to the different neurons recognized.

One of the distinctive characteristics of the SmRG is its ability to trace the axon topology of OP fibers while maintaining 3D volumetric information on neurons and their arbours. Indeed, the structure obtained with the SmRG is a smooth three-dimensional volume with voxel-resolution details on neuron morphology; a feature not available from swc structures. As a consequence, the SmRG reconstructions in Figure 7 appear thicker than the 3D rendering of *.swc gold-standards.

**Fig 7.**
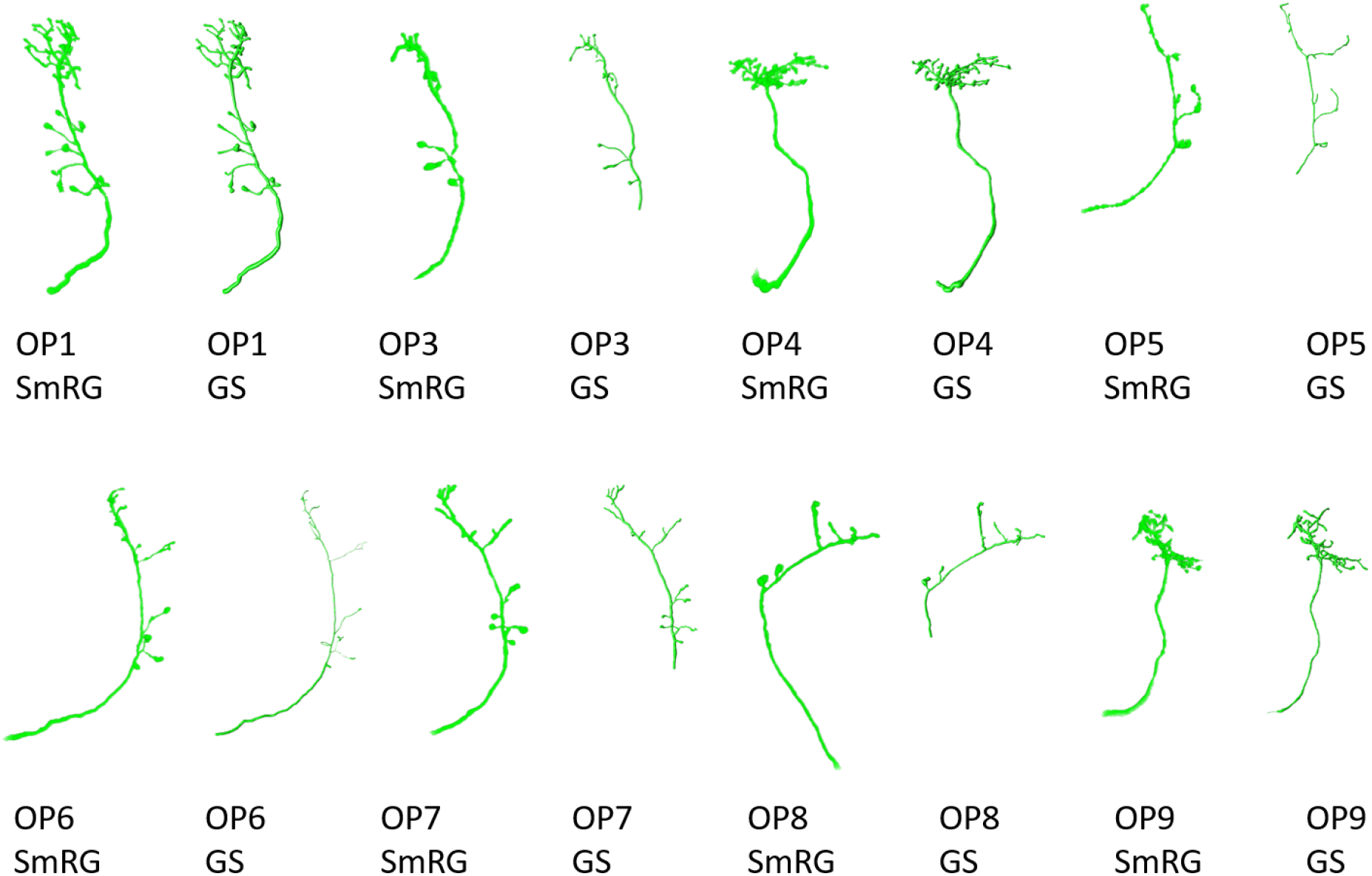
OP fibers segmented with SmRG and compared with the DIADEM gold-standard (GS). Please note that for OP3, OP5, OP7 and OP8 the gold standard reconstruction misses some terminal branches (see DIADEM FAQ at http://diademchallenge.org/faq.html)

As evident from the figure, some terminal branches of OP fibers are not comprised in the manually traced gold standard, since they have no effect on DIADEM metrics (Brown et al. 2011; Gillette et al. 2011). Nonetheless, the SD, SSD and SSD% metrics used in this work are naturally biased by these missing branches. Thus, the comparison between automatic reconstructions and gold standard were limited to those branches included in by the DIADEM gold standard.

When evaluated against other SoA tools, the SmRG was observed to be comparable in terms of SD. On the other hand, our algorithm achieved the lowest values of SSD among all tools considered (with the exception of segmentation of OP5). It should be noted that the value of SSD% was higher for the SmRG with respect to other algorithms, since the estimation of the skeleton from the 3D output of SmRG produced a higher number of nodes compared to the other methods (Fig 8).

**Fig 8.**
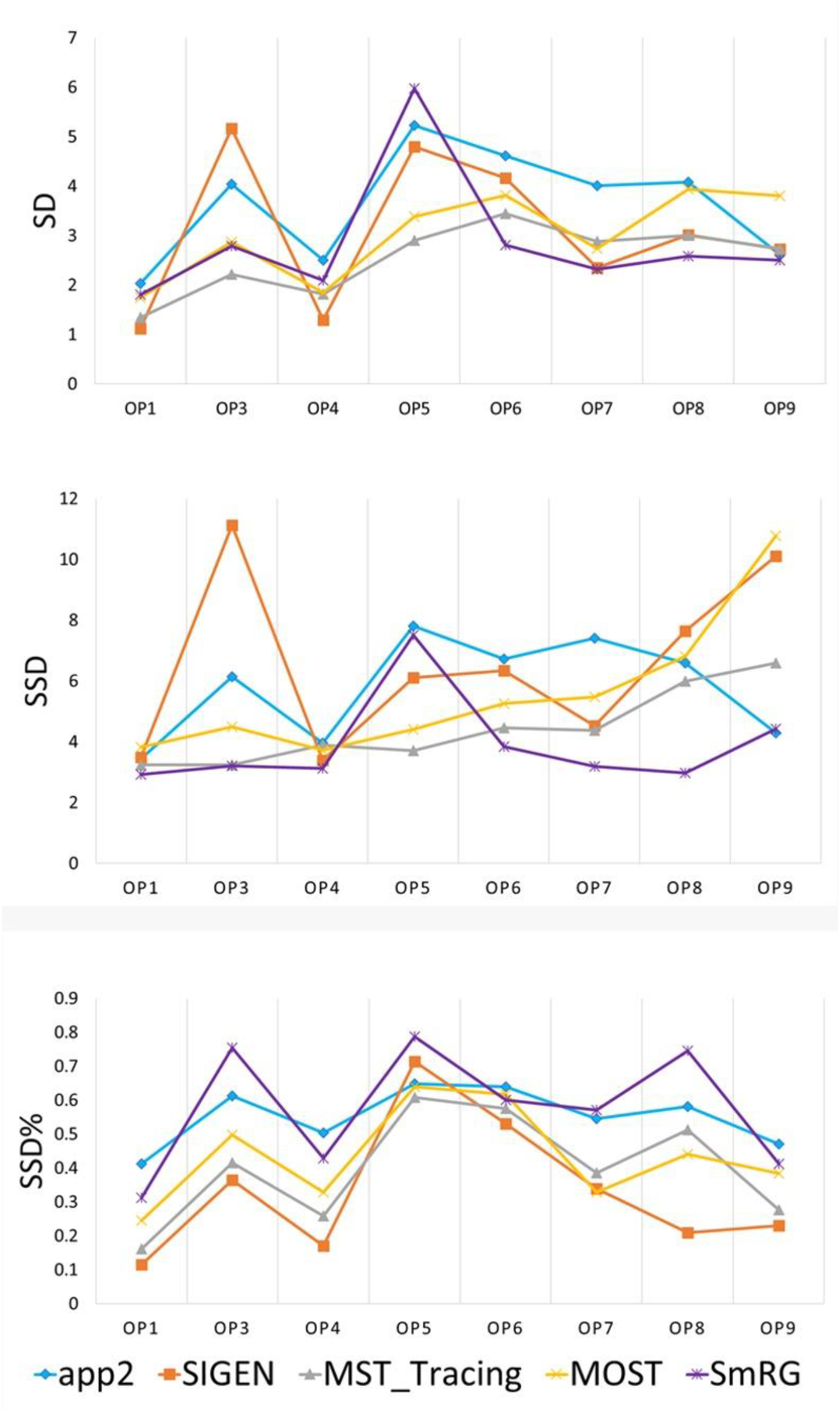
Accuracy of SmRG and SoA tools against the DIADEM gold standard for different OP fibers. A) SD B) SSD and C) percentage SSD.

In Fig 9 the average precision, recall and F-score across OP fibers are reported for SmRG and SoA tools as a function of the value of S. For S=5, the SmRG outperforms other tools in terms of F-score which highlights its ability to segment OP fibers with high accuracy.

**Fig 9.**
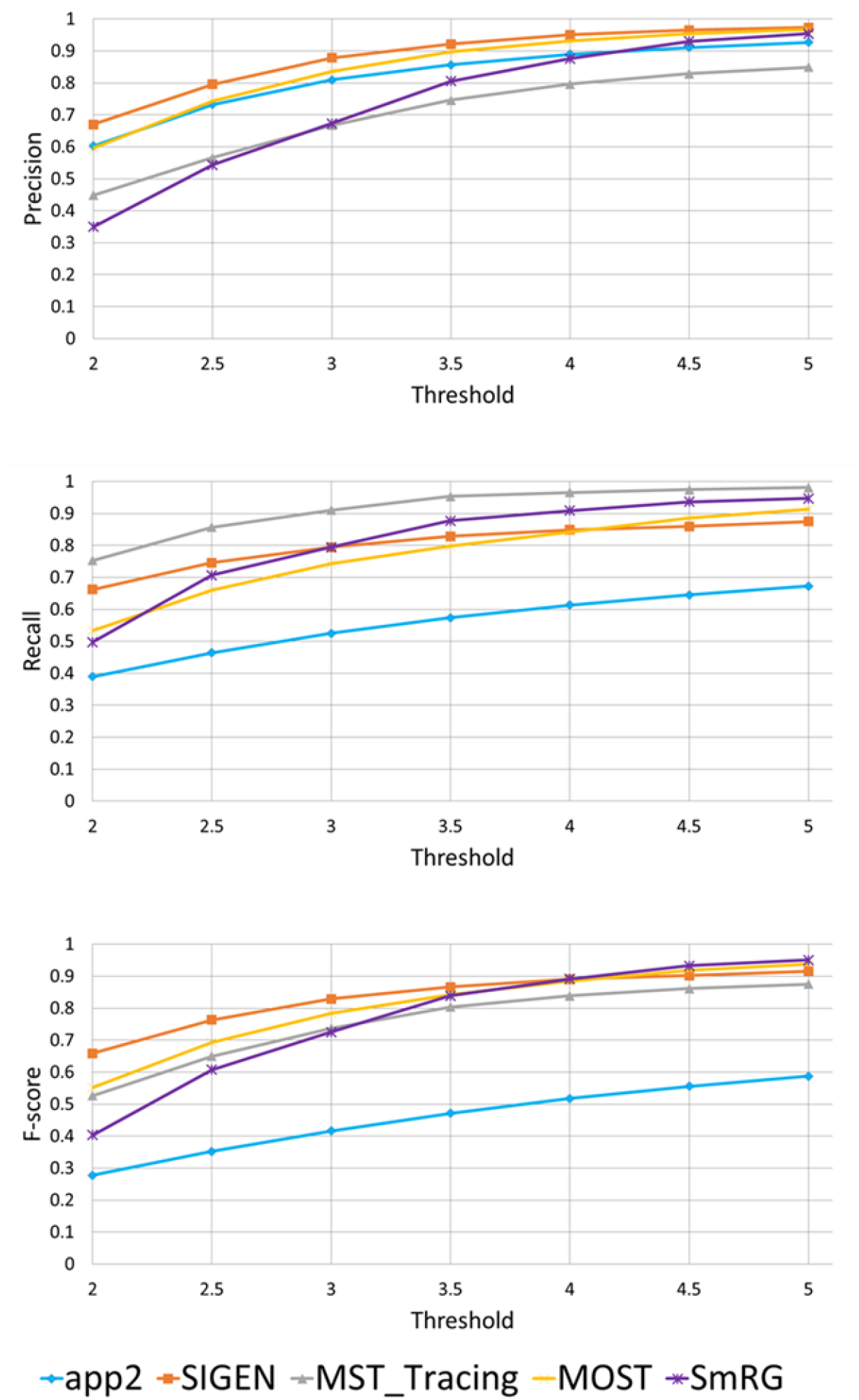
Precision, Recall and F-Score for varying thresholds of SSD evaluation. SmRG and SoA tools have similar performance for increasing values of the threshold. For thresholds greater than 4 voxels, SmRG has the highest F-Score. For S=5, we obtained P = 0.9538 ± 0.0350, R = 0.9770 ± 0.0183 and F=0.9651 ± 0.0248 (mean ± st. deviation) for the SmRG.

## 5 DISCUSSION

The SmRG for the automatic segmentation of microscopic data exploits the signal statistics typical of confocal and 2-photon images (Calapez and Rosa 2010). Datasets representing neural tissues from different species, processed using different protocols (i.e. clarified murine cerebella and Drosophila brains fixed using classical procedures) and acquired with different imaging tools (i.e. confocal and two photon microscopy) were used to test the algorithm. The goodness of the SmRG reconstruction was compared with manually traced gold-standards as well as with algorithms available in the SoA.

A quantitative analysis of the SmRG’s accuracy with PC datasets was performed for 3 different neurons, whose manually segmented counterpart was available in Magliaro et al., 2017 ^35^. Although a limited set of neurons were analysed, the reconstructions of the SmRG and the manually-segmented gold standards were comparable; moreover, the seeding and RG procedure was shown to be robust and independent of initial conditions. The analysis performed on PCs from clarified tissues highlighted the efficacy of the algorithm developed in isolating single neurons from densely-packed data with respect to some of the most widely used single neuron reconstruction tools available in the SoA (i.e. app2, MOST, MST-tracing, SIGEN) (Basu and Racoceanu 2014; Ikeno et al. 2018; Ming et al. 2013; Xiao and Peng 2013). In particular, none of the SoA tools allowed the reconstruction of 3D neuron morphology from the confocal stacks representing neurons in their native 3D context, limiting the evaluation of the SmRG’s performance to a visual comparison. Indeed, many of the SoA algorithms perform extraordinarily well with low-quality images possessing noisy points, large gaps between neurites and non-smooth surfaces (Liu et al. 2016), since they were likely developed specifically for such purposes. On the contrary, they may perform modestly or even fail in reconstructing densely-packed neurons(Hernandez et al. 2018), such as PCs in the murine cerebella because the images have very different properties (i.e. a large number of pixels with high intensities). As it was specifically developed for clarified brain tissue, the SmRG could provide a valid alternative to SoA tools for the segmentation of such datasets. Further validation of the algorithm on other tissues is currently ongoing to confirm its robustness when dealing with different neuronal types.

Reconstructions of OP fibers from the DIADEM challenge resulted in a comparable performance between the SmRG and well-established tools for neuron reconstruction in terms of SD, SSD and SSD%. Specifically, the algorithm proposed here outperformed other tools in terms of SSD, which quantifies the discrepancy between two outcomes (Peng, Long, and Myers 2011), in almost all reconstructions. On the other hand, the SmRG exhibited higher values in the SSD% score. It should be noted that the skeleton of SmRG reconstructions is derived from a thinning procedure of volumetric datasets, which results in a higher number of nodes with respect to the DIADEM ones. This may bias the SSD% towards higher values (Liu et al. 2016).

The precision and recall of SmRG outcomes with respect to the manually traced gold-standard provided by the DIADEM highlighted the exceptional performance of our tool (comparable to SoA algorithms in the segmentation of OP fibers). In particular, for the highest values considered of the tolerance threshold, the SmRG’s average values of precision, recall and f-score were all above 95%. This suggests that, although the algorithm was developed for segmenting neurons from clarified cerebral tissue, segmentation procedures based on local signal and noise statistics may be a successful strategy for delivering an adaptive and generalised algorithm, applicable to different contexts.

We would like to underline that the SmRG was not compared with SoA segmentation approaches in terms of computational times. Indeed, tools such as app2, MST, SIGEN and MOST outperform our algorithm as they provide faster segmentations with comparable accuracy and precision (Fig 8) for sparsely labeled data, while they fail when performing segmentations of densely-packed neurons. The SmRG was, in fact, specifically aimed at addressing the lack of tools able to deal with the intricacies of mammalian brains.

Finally, when two neurons naturally touch each other and the signal intensity is high, SmRG may reconstruct the two objects as one, thus requiring their post-splitting. An ad hoc watershed-based routine for separating neurons is provided at http://www.centropiaggio.unipi.it/smrg-algorithm-smart-region-growing-3d-neuron-segmentation. Further improvements are ongoing to optimize the tool and its computational time.

## 5. CONCLUSION

Despite the numerous attempts addressed at 3D neuron reconstruction, little attention has been paid to delivering automatic and robust methods capable of dealing with the large variability of datasets representing densely-packed neurons, as well as for digitizing the morphology and volumetric characteristics of the segmented structures. As a result, the majority of algorithms are only able to handle with sparsely labelled data, compelling neuroscientists to manually segment images representing intricate neuronal arborisations and to reducing 3D space-filling neurons to skeletonised representations.

The SmRG, an open-source Matlab-based algorithm for the segmentation of complex structures in 3D confocal or 2-photon image stacks overcome these setbacks. It provides an accurate reconstruction of 3D neuronal morphology acquired using confocal microscopy, which accounts for 80% of user needs in imaging facilities. The SmRG can potentially be extended to other imaging modalities (e.g., super-resolution microscopy) adopting the same statistical framework for identifying the signal and noise distribution from 3D images.

In addition, our tool allows the extraction of several useful morphological features from the segmented neurons. Preserving the volumetric information is an essential step for deciphering the Connectome. Besides structural mapping, from a biological perspective, digital 3D neuron reconstruction is crucial for the quantitative characterization of cell type by morphology and the correlation between morphometric features and genes (e.g. between wild-type and model animals) or patho-physiology (e.g. the detection of neuronal morphological anomalies in diseased individuals compared to healthy ones) (Acciai et al. 2016). In conclusion, the SmRG can facilitate the identification of the different neural types populating the brain, providing an unprecedented set of morphological information and new impetus towards connectomic mapping.

## Acknowledgments

The authors would like to thank prof. Vincenzo Positano for the useful suggestions during the first steps of the algorithm design and Gabriele Paolini e Gianluca Rho for their help in SmRG implementation. The authors would also like to thank Michele Scipioni for useful suggestions during the development of the EM algorithm. CM is thankful to Fondazione Veronesi for her Post-Doctoral Felloshwip.

## Author Contributions

ALC, CM and NV: conception and design of the segmentation tool; ALC: implementation and testing of the algorithm; ALC, CM, AA and NV: interpretation of data and drafting of the manuscript.

## Appendix 1 The Mixture Model

In its original version (Calapez and Rosa, 2010), the model is supposed to describe K different fluorescence levels or classes; the k-th class is described by the linear mixture model:

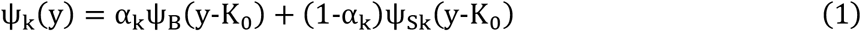

where y, K_0_ and α_k_ denote the pixel intensity level, the system offset and the mixture parameter respectively. ψ_B_ is the distribution for the background pixels and is modeled according to a discrete normal distribution, with variance v_B_ and mean K_0_, and ψ_Sk_ is the intensity distribution of the k-th class pixels, described by a negative-binomial distribution with variance v_Sk_ and mean μ_Sk_. In accordance with (Calapez and Rosa, 2010) the negative-binomial distribution is re-parameterized in terms of

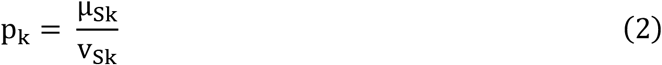

and

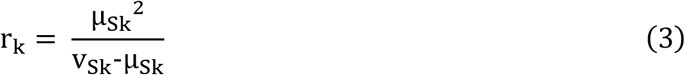

For region growing purposes, it is reasonable to assume the presence of a single class k of pixels, at least locally. In this case, the complete model for a pixel y_l_ is described by the 5-parameter distribution

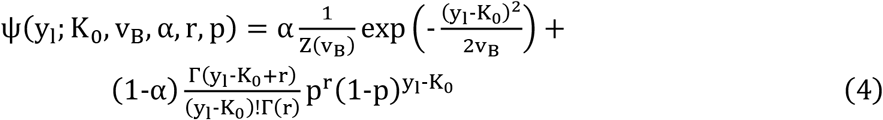

where all the parameters are real valued except for K_0_ which is an integer and α ∈ [0,1].

The model fitting is done by means of an Expectation-Maximization (EM) algorithm in which:

1. p and r are obtained by the method of moments (equations 2 and 3)
2. K_0_ and v_B_ are given by the maximization of the log-likelihood

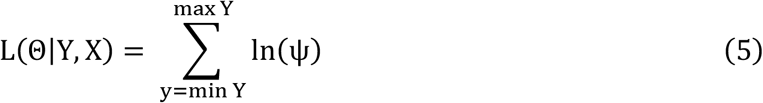
3. α is given by the posterior density

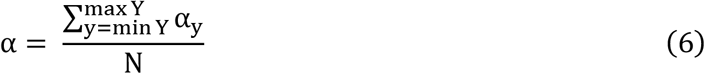

